# CslA and GlxA from *Streptomyces lividans* form a functional cellulose synthase complex

**DOI:** 10.1101/2023.11.20.567928

**Authors:** Xiaobo Zhong, Simone Nicolardi, Ruochen Ouyang, Manfred Wuhrer, Chao Du, Gilles van Wezel, Erik Vijgenboom, Ariane Briegel, Dennis Claessen

## Abstract

Filamentous growth of streptomycetes coincides with the synthesis and deposition of an uncharacterized protective glucan at hyphal tips. Synthesis of this glucan depends on the integral membrane protein CslA and the radical copper oxidase GlxA, which are part of a presumably large multiprotein complex operating at growing tips. Here, we show that CslA and GlxA interact by forming a protein complex that is sufficient to synthesize cellulose in vitro. Mass spectrometry analysis revealed that the purified complex produces cellulose chains with a degree of polymerization of at least 80 residues. Truncation analyses demonstrated that the removal of a significant extracellular segment of GlxA had no impact on complex formation, but significantly diminished activity of CslA. Altogether, our work demonstrates that CslA and GlxA form the active core of the cellulose synthase complex and provides molecular insights into a unique cellulose biosynthesis system that is conserved in streptomycetes.

**Significance:** Cellulose stands out as the most abundant polysaccharide on Earth. While the synthesis of this polysaccharide has been extensively studied in plants and Gram-negative bacteria, the mechanisms in Gram-positive bacteria have remained largely unknown. Our research unveils a novel cellulose synthase complex formed by the interaction between the cellulose synthase-like protein CslA and the radical copper oxidase GlxA from *Streptomyces lividans*, a soil-dwelling Gram-positive bacterium. This discovery provides molecular insights into the distinctive cellulose biosynthesis machineries. Beyond expanding our understanding of cellulose biosynthesis, this study also opens avenues for exploring biotechnological applications and ecological roles of cellulose in Gram-positive bacteria, thereby contributing to the broader field of microbial cellulose biosynthesis and biofilm research.

## Introduction

Cellulose, a linear β-1,4-glucan, is the most abundant natural biopolymer on our planet and has great potential for applications in biotechnology and biofuel production (1). Cellulose is found predominantly in green plants, where it is used as a structural component of the cell wall (2). However, it is also abundantly found as a component of the extracellular matrix of bacterial biofilms. The biosynthesis of cellulose is executed by sophisticated multiprotein complexes (3). A key component of these complexes is membrane-integrated processive synthases, which belong to the glycosyltransferase (GT) −2 superfamily. Members of this family also include synthases for chitin and alginate with β-1,4-linkages, and hyaluronan with β-1,3- and 1,4-linkages (4). In plants, cellulose is synthesized by cellulose synthases (CesAs), which assemble into a large six-lobed rosette-shaped complex that assembles the extruded glucan chains into microfibrils (5, 6). Homologues of CesA, designated as cellulose synthase-like (Csl) proteins, can synthesize other polysaccharides including xyloglucan (7), glucomannan (8) and β-1,3:1,4-glucan (9). In Gram-negative bacteria, cellulose synthesis is performed by the so-called BcsA-BcsB complex (10). BcsA is an integral membrane protein carrying the cytosolic catalytic domain. BcsB is a periplasmic protein that interacts with BcsA. This interaction is mediated via transmembrane helices in both proteins and probably contributes to the unidirectional movement of the glucan chain during translocation (10). The synthesis of cellulose by the combined action of the synthase BcsA and the accessory protein BcsB is commonly found in Gram-negative proteobacteria, like *E. coli* K-12 (11), *Rhodobacter sphaeroides* (10) and *Gluconacetobacter xylinus* (12). Recently, a functional homolog of the BcsA-BcsB complex was found in the Gram-positive bacterium *Clostridioides difficile* (13). While the systems in Gram-negative bacteria have been characterized, little is known about the systems in Gram-positive bacteria.

The Gram-positive streptomycetes possess a cellulose synthase-like protein, called CslA (14, 15). As saprophytes, streptomycetes are ubiquitous in soil environments. They are filamentous, multicellular bacteria that form reproductive aerial structures after a period of vegetative growth. Like BcsA, CslA belongs to the glycosyl transferase GT-2 superfamily, and is required for glucan synthesis at hyphal tips (14). Following synthesis, the glucan becomes exposed on the surface of the multi-layered cell wall (16). The gene encoding CslA is contained in a conserved gene cluster in most streptomycetes. Two genes in this cluster, *lpmP* and *cslZ*, were proposed to mediate the secretion of the produced glucan through the peptidoglycan layer (17). *cslA* is transcriptionally and translationally coupled to *glxA*, encoding a radical copper oxidase (18). Mature GlxA coordinates a copper radical through the covalent cross-linking of Cys and Tyr residues, termed the Tyr-Cys redox cofactor (18–20). Interestingly, the deletion of either *cslA* or *glxA* genes blocked the deposition of the glucan at hyphal tips (14, 18), implying that both proteins are crucial for synthesis. However, the structure of GlxA shares no structural and functional homology with the auxiliary BcsB proteins that co-operate with BcsA (-like) proteins, making the CslA-GlxA system novel. Whether CslA and GlxA form a functional complex was unknown.

In this study, we heterologously expressed CslA and GlxA in *E. coli* and found that these proteins form a functional complex *in vitro* that can synthesize cellulose. More specifically, enzymatic degradation of the polymer combined with MALDI FT-ICR MS analysis indicated that the CslA-GlxA complex produces cellulose chains of at least 60-80 glucose units with UDP-glucose as substrate. Altogether, our work unambiguously demonstrates the potential for cellulose biosynthesis in a Gram-positive bacterium for the first time.

## Results

### Streptomyces CslAs belong to a distinct group of cellulose synthases

CslA is crucial for synthesizing a glucan at hyphal tips in *Streptomyces*, which is required for morphogenesis(14, 18). A phylogenetic comparison between cellulose synthases showed that CslA from the streptomycetes (*S. avermitilis*, *S. clavuligerus*, *S. coelicolor, S. griseus*, *S. lividans, S. scabies* and *S. venezuelae*) comprise a distinct group of synthases. These are derived from a common ancestor shared with the bacterial BcsA-family of synthases and the plant-based CslA and CslC synthases (Fig. 1). The latter proteins synthesize glucomannan consisting of β-1,4-linked alternated mannose/glucose units (21) and xyloglucan consisting of a β-1,4-linked glucan backbone (22), respectively. The common feature of bacterial BcsA (23) and plant CslA (21)/CslC (22) is that they can both utilize UDP-Glc to either synthesize cellulose or a hybrid polysaccharide. The alignment of amino acid sequences of CslA from *S. lividans* and the well-characterized BcsA from *Rhodobacter spaeroides* revealed that these two proteins share 27% sequence identity and CslA putatively possesses several highly conserved motifs from bacterial cellulose synthases, including the UDP-coordinating sites (DXD and TED) (10), a putative gating loop (FXVTXK) (24), the glucan terminal disaccharide binding sites (QXXRW) (10), the Mg^2+^-binding histidine residue adjacent to DXD (10), and also elements of the c-di-GMP binding PilZ domain (24) (Fig. S1). In view of the high sequence similarity and common ancestor with bacterial BcsAs, we speculate that *Streptomyces* CslAs can utilize UDP-Glc as substrate.

**Figure 1.**
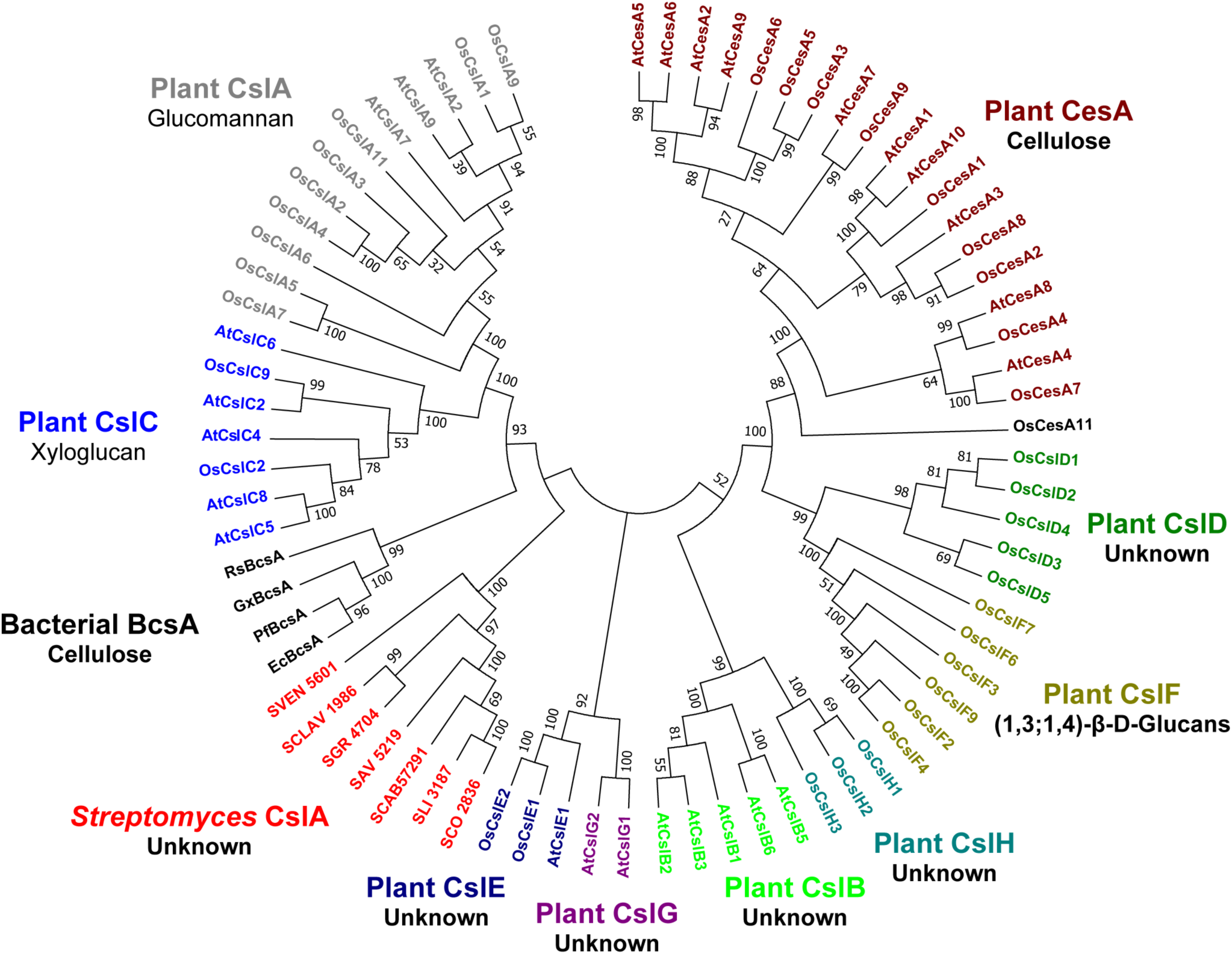
Phylogenetic analysis of cellulose synthase or synthase-like proteins from plants and bacteria. The analysis was done using bacterial cellulose synthases (BcsAs) from *E. coli* K-12, *Komagataeibacter xylinus* and *Pseudomonas fluorescens*, plant cellulose synthases (CesAs) and synthase-like (Csl-) proteins CslA/B/C/D/E/F/G/Hs from *Arabidopsis thaliana* and *Oryza sativa*, and streptomycetes cellulose synthase-like CslAs from *S. coelicolor* (SCO2836), *S. lividans* (SLI3187), *S. venezuelae* ATCC10712 (SVEN_5061), *S. clavuligerus* ATCC 27064 (SCLAV1986), *S. griseus* (SGR4704), *S. avermitilis* MA-4680 (SAV5219), and *S. scabies* (SCAB57291). The phylogenetic tree was built using MEGA7 with the Maximum likelihood method, of which the bootstrap value was inferred from 100 replicates. Proteins from distinct groups are colored differently and the corresponding products are indicted underneath.

### Membrane topology analysis of GlxA

The *cslA* and *glxA* genes are located in the same operon and function co-operatively in glucan synthesis (17). Previous structure predictions indicated that CslA has seven transmembrane helices, while GlxA was predicted to possess a transmembrane helix and a short N-terminal membrane anchor (18). To test if CslA and GlxA form a membrane-associated complex, we first determined the membrane topology of GlxA. Computational analyses indicate that GlxA is composed of a cytoplasmic motif (aa. 1-12), a transmembrane helix (aa. 13-32), and an extracellular domain (aa. 33-645) (Fig. S2). Additionally, a weak signal peptide is situated at the N-terminus with a predicted cleavage site at G25 (Fig. S2). To confirm the presence of the signal peptide responsible for GxlA secretion, a β-lactamase assay was employed, wherein the enzyme was fused to the C-terminus of GxlA (for detailed methodology, see the Materials and Methods section). Significantly, *E. coli* cells expressing GlxA_1-32_-BlaM_NS_ and GlxA_FL_-BlaM_NS_ fusions exhibited resistance to ampicillin, comparable to the positive control strain expressing β-lactamase with its native signal peptide (BlaM_FL_) (Fig. 2A). The findings establish that GlxA’s N-terminal residues 1-32 can facilitate the secretion of β-lactamase into the periplasm. This provides evidence for the presence of a signal peptide within this region, which also encompasses a transmembrane (TM) anchor responsible for anchoring the protein to the cell membrane. Indeed, GlxA was previously shown to associate with the membrane via non-covalent interactions (18). Thus, the predominant portion of GlxA, encompassing its central propeller-shaped Domain 1 (aa. 55-142 and 261-541 for Region-1 and Region-2 respectively), the extended Domain 2 (aa. 143-260), and Domain 3 (aa. 542-645), resides in the extracellular space (Fig. 2D).

**Figure 2.**
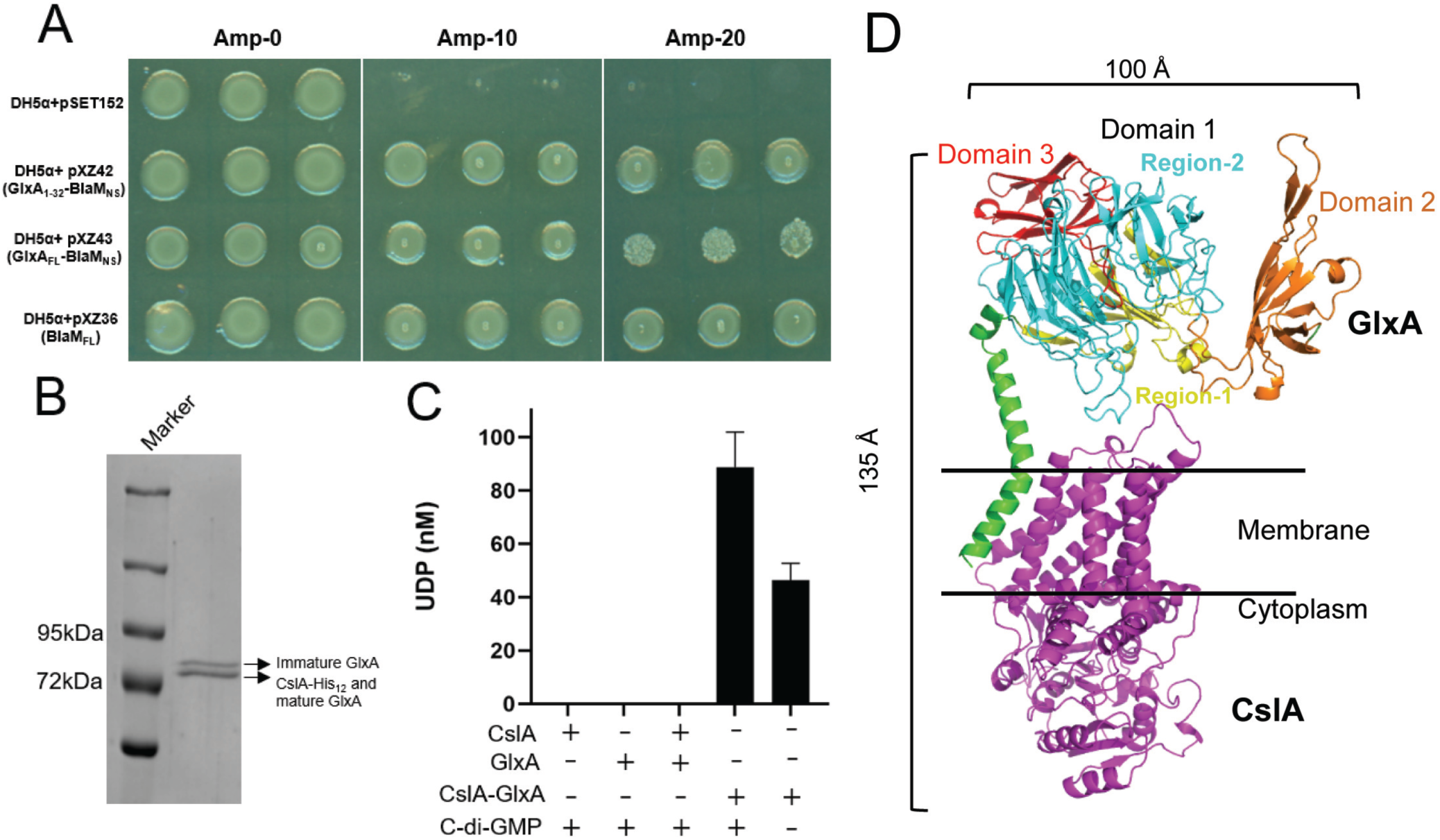
CslA and GlxA are required for cooperative glucan synthesis *in vitro*. (**A**) Membrane topology analysis of GlxA. DH5α containing plasmids pSET152 (empty), pXZ42 (expressing the β-lactamase in which its original N-terminal signal peptide is replaced by aa. 1-32 of GlxA; GlxA_1-32_-BlaM_NS_), pXZ43 (expressing the β-lactamase without its original signal peptide fused to the C-terminus of full length GlxA; GlxA_FL_-BlaM_NS_) or pXZ36 (the β-lactamase containing its original signal peptide) were spotted on plates with 0, 10 or 20 μg ml^-1^ ampicillin. Plates were imaged after overnight growth at 37 °C. (**B**) Purification of the CslA-GlxA complex expressed in *E. coli* BL21 (DE3). SDS-PAGE of the affinity and size-exclusion chromatography purified CslA-GlxA complex stained with Coomassie Brilliant Blue. Arrows indicate which protein is identified at that position by immunoblotting. The upper band indicates immature GlxA (lacking the Tyr-Cys cofactor), while the lower band indicates CslA-His_12_ overlapping with mature GlxA (containing the Tyr-Cys cofactor). (**C**) GlxA is required for the catalytic activity of CslA. Reactions were performed at 37°C for 60 min by incubating 0.1 mg ml^-1^ of the individual monomers, a mixture of monomers or the co-purified complex using 5 mM UDP-Glc, 5 mM cellobiose, 20 mM MgCl_2_ and 30 μM c-di-GMP. Additionally, the activity of the co-purified complex was assessed separately without the addition of c-di-GMP. The catalytic activity of enzymes was quantified by measuring free UDP with the UDP-Glo^TM^ glycosyltransferase assay. Every measurement was performed in triplicate. Error bars represent the standard error of the mean. (**D**) AlphaFold 2.0 prediction of the CslA-GlxA complex. The overall dimensions of the complex are indicated by size bars (top and left).

### ClsA and GlxA form a complex

To test if CslA and GlxA form a complex, we co-expressed both proteins in *E. coli* and performed pull-down studies. To this end, full-length CslA was expressed with a 12-Histidine tag at its C-terminus, while non-tagged GlxA has aa. 1-11 replaced with the PelB signal sequence (for efficient translocation into the periplasm, Fig. S2), while retaining GlxA’s N-terminal transmembrane helix (TMH, Fig. S2). Following co-expression, CslA-His_12_ was purified using affinity chromatography and used to pull-down non-tagged GlxA in the co-expression studies. Meanwhile, we also purified CslA-His_6_ and GlxA-His_6_ (see the Materials and Methods section), which were used as controls. Interestingly, SDS-PAGE analysis of the pull-down samples indicated that the single peak detected with size exclusion chromatography was in fact a mixture of two proteins with masses of 73.1 and 69.5 kDa (Fig. 2B, Fig. S3A). Western analysis confirmed that CslA successfully pulled-down GlxA (Fig. S3B), which demonstrated that CslA interacts with GlxA and formed a complex in *E. coli*. Immunoblotting with specific antibodies revealed one band for CslA and two bands for GlxA in the CslA-GlxA complex (Fig. S3B). This migration pattern of GlxA corresponds to the presence (lower band) and absence (upper band) of the Tyr-Cys redox cofactor (19, 20). This observation suggests that both the mature and immature forms of GlxA can be present within the CslA-GlxA complex.

To predict the regions of GlxA required for forming the complex with CslA, AlphaFold 2.0 was used, which predicted that CslA and GlxA form a heterodimer (Fig. 2D). The N-terminal TMH of GlxA is hypothesized to interact with the transmembrane regions of CslA in the membrane through hydrophobic interactions. To corroborate these *in silico* analyses, we generated two truncated variants of GlxA, both tagged with a C-terminal Flag-tag: GlxA^12-261^, lacking Region-2 and Domain 3, and GlxA^12-54^ encompassing only the N-terminal TMH. Subsequently, these truncated GlxA proteins were used in pull-down assays with CslA-His_12_ (Fig. S2 and S4A). Immunoblotting analyses confirmed that CslA-His_12_ effectively precipitated Flag-tagged GlxA^12-261^ (30.4 kDa), as detected by both by α-GlxA and α-Flag antibodies (Fig. S4B). However, CslA-His_12_ did not interact with Flag-tagged GlxA^12-54^ (7.8 KDa) (Fig. S4B). Collectively, these results underscore that, apart from the TMH, other segments of GlxA play a crucial role in its interaction with CslA. Altogether, these analyses demonstrate that CslA and GlxA form a binary complex on the cell membrane.

### Synthesis of cellulose by the CslA-GlxA complex

The homology of CslA to cellulose synthases prompted us to investigate if the CslA-GlxA complex could synthesize cellulose. In cellulose polymerization reactions, the catalytic domains of the cellulose synthases transfer glucose units from the donor substrate UDP-Glucose (Glc) to the non-reducing ends of nascent cellulose chains, thereby releasing UDP (25). In agreement, UDP release was detected when the substrate was incubated with the CslA-GlxA complex (Fig. 2C). No UDP release was detected when CslA and GlxA were used individually or when they were mixed after purifying them separately (Fig. 2C). Together, this demonstrates that the tight integration of GlxA in the complex is essential for UDP-glucose turnover. Remarkably, it was observed that the CslA-GlxA complex exhibited UDP-glucose turnover even in the absence of c-di-GMP (Fig. 2C). However, its maximum activity was achieved in the presence of this activator molecule, with optimal performance observed at a substrate concentration of 6.8 mM and a temperature of 38°C (Fig. S5). Furthermore, the CslA-GlxA^12-261^ complex exhibited a significantly reduced activity, which was approximately 40% of that observed in the wild-type CslA-GlxA complex (Fig. S4C). This strongly implies that the presence of the intact domain 1 of GlxA is required for realizing the complex’s full catalytic potential.

To establish that the complex is responsible for the production of cellulose, we purified the insoluble product and incubated it with a cellulase or a β-1,3-specific glucanase (Fig. 3A). Contrary to cellulase, the β-1,3-specific glucanase was unable to degrade the *in vitro* produced glucan (Fig. 3A), consistent with it being a β-1,4-glucan. Indeed, mass spectrometric analysis of the products released after degradation with cellulase revealed a hexose ladder that was highly similar to the enzymatic degradation products generated from commercial cellulose (Fig. 3B, Fig. S6). In both cases, polydisperse oligo- and poly-glucose carbohydrates (in the range of ± 6-86 glucose residues) could be identified. CID MS/MS analysis of selected precursor ions, corresponding to hexa-glucose oligosaccharides (i.e. Glc_6_; detected at *m/z* 1013.3168) obtained from cellulase digestion of the *in vitro* produced β-1,4-glucan as well as commercial cellulose resulted in virtually identical fragmentation spectra (Fig. S7), further corroborating that the polymer synthesized by the CslA-GlxA complex is cellulose.

**Figure 3.**
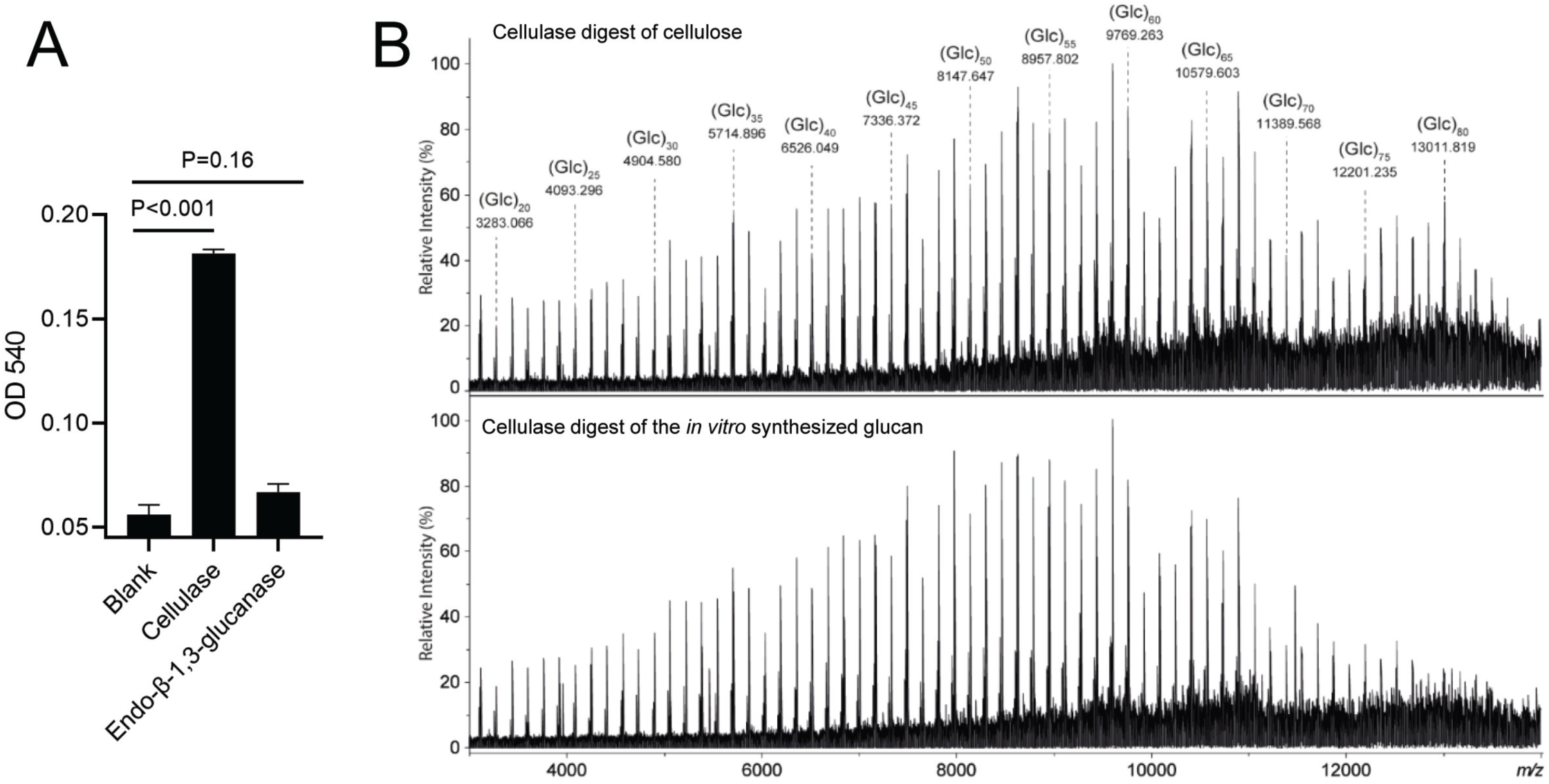
Characterization of glucans produced by CslA-GlxA *in vitro*. Synthesis of glucans was performed in 1 ml reactions containing 0.5 mM CslA-GlxA, 5 mM cellobiose, 6.5 mM UDP-Glc, 20 mM MgCl_2_ and 30 μM cyclic-di-GMP. After incubation for 60 min at 37°C, the glucans were collected by centrifugation, washed, and reduced sugars were quantified by DNS reagent after enzymatic degradation (**A**) and MALDI-CID FT-ICR MS analysis (**B**). For the enzymatic degradation, 20 μg ml^-1^ cellulase and 20 μg ml^-1^ β-1,3-glucanase were used and the absorbance at a wavelength of 540 nm (OD540) was used to evaluate polymer degradation. Reactions were done in triplicate and error bars indicate the standard error of the mean. (**B**) Enlargement of MALDI FT-ICR mass spectra in the *m/z*-range of 2000-14000. Mass spectra were obtained from the analysis of the cellulase digest of cellulose (top panel) and the cellulase digest of the *in vitro* synthesized glucan (bottom panel).

## Discussion

Streptomycetes are prolific antibiotic-producing Gram-positive bacteria that grow by tip extension. The continuous remodeling of the cell wall associated with tip growth makes these sites relatively vulnerable. To provide protection, streptomycetes produce an apical glucan that covers the peptidoglycan layer. Synthesis and deposition of this glucan is exerted by a multiprotein complex operating at growing tips. Here we show that CslA, together with its accessory protein GlxA, are capable and sufficient for glucan synthesis. Furthermore, we show that these proteins form a binary complex in *E. coli* that can synthesize cellulose *in vitro*. Together, this work provides new insights into a unique cellulose biosynthesis system in a multicellular Gram-positive bacterium.

Cellulose is the most abundant organic compound on Earth and is produced by a wide range of organisms. In bacteria, the synthesis of cellulose is best characterized in Gram-negative via the BcsA-BcsB complex. Here, a pronounced interaction between BcsA and BcsB is mediated via BcsB’s C-terminal TM anchor, which contributes to the stabilization of the glucan secretion channel formed in TM helices of BcsA (10). Like in BcsA-BcsB, cellulose synthesis in *Streptomyces* is also accomplished by a dual-component system comprising the cellulose synthase CslA and the radical copper oxidase GlxA. Interestingly, CslA and BcsA share 27% sequence identity (Fig. S1), but no homology is present between GlxA and BcsB. AlphaFold predicted that GlxA contains a TM helix, which would firmly anchor GlxA into the membrane and form a complex with CslA (Fig. 2B). This suggests that GlxA’s TM anchor may fulfill a comparable role in the stabilization of the complex like BcsB’s TM anchor in the BcsA-BcsB system. In the BcsA-BcsB system, the presence of the N-terminal TM anchor BcsB alone is adequate for the formation of a functional BcsA-BcsB complex (23). However, in contrast, the formation of a fully functional CslA-GlxA complex requires both the N-terminal TM anchor and the intact extracellular regions of GlxA to be present (Fig. S4).

Interestingly, full activity of the ClsA-GlxA complex also requires the presence of c-di-GMP, a second messenger commonly required for synthesis of extracellular polysaccharides (EPS). Binding of c-di-GMP relies on specific motifs within the synthases: the PilZ domain (RXXXR and D/NXSXXG motif) and GGDEF I site (RXXD motif) (26). CslA exhibits a partial PilZ domain characterized by the presence of the RXXXR motif (see Fig. S1). As the catalytic activity of the CslA-GlxA complex in the absence of c-di-GMP is comparable to the CslA-GlxA^12-261^ complex (Fig. 2C, Fig. S4), we hypothesize that c-di-GMP binding may not only involve CslA, but also GlxA region-2. This observation raises the intriguing possibility of a novel, fully functional c-di-GMP binding domain potentially emerging upon the formation of the CslA and GlxA complex. Indeed, binding of c-di-GMP by BldD, a master regulator controlling the switch from vegetative growth to sporulation, depends on a unique RXD-X8-RXXD signature as has been reported in streptomycetes (27, 28). The detailed interactions between CslA and GlxA and the identification of the c-di-GMP binding site(s) need to be further investigated in the future.

In addition to *cslA* and *glxA*, biosynthesis of *Streptomyces* cellulose also involves *cslZ*, which lies downstream of *cslA-glxA* operon (17). Interestingly, the *cslA-glxA-cslZ* gene cluster of the *Streptomyces* cellulose synthesis system is highly similar to the organization of *bcsA-bcsB-bcsZ* in Gram-negative bacterial cellulose systems, but also in a recently described system in the Gram-positive bacterium *Clostridium* (13). The common feature between all these systems is that synthesis of the glucose chain is carried out by a core synthase (CslA, CcsA or BcsA) and a second partner (GlxA, CcsB or BcsB) lacking homology. Nevertheless, secretion to the outer surface is accomplished by distinct mechanisms, in which Gram-negatives use an outer membrane pore formed by BcsC (29). Instead, streptomycetes use two designated proteins, LpmP and CslZ, to create a passage through the thick PG layer (17).

While our work unambiguously shows that CslA-GlxA can synthesize cellulose *in vitro*, it remains to be established whether this is also the glucan that is produced *in vivo*. The glucanase CslZ shows hydrolytic activity on different substrates including PG, various types of cellulose but also α-chitin (17). Instead, BcsZ was only found to be active on carboxymethyl-cellulose (30). The promiscuous activity of CslZ perhaps suggests that CslA is also able to utilize different sugars based on their availability. In fact, this promiscuity has been reported before for a cellulose synthase-like protein from *Arabidopsis*, which can incorporate both mannose or glucose into large-linked homo- or heteropolymers (21). The ability to incorporate different sugars could provide a tremendous benefit for streptomycetes in nature, which face a wide variety of substrates in nature. Our isolation of a minimal, yet functional CslA-GlxA complex provides a powerful tool to study this possibility and draw an outline of many crucial steps during cellulose microfibril formation in streptomycetes and their roles in providing hyphal tip protection.

## Materials and Methods

### Plasmid construction for heterologous production of CslA and GlxA in E. coli

All primers and plasmids used in this article are listed in Tables S1 and S2, respectively. The *cslA* gene (SLI3187) was amplified from genomic DNA of *Streptomyces lividans* using primers 3187-F and 3187-R. The amplified product was purified and ligated as a NdeI/HindIII fragment into Pet21a (a gift from Dr. Jonathan A. R. Worrall, University of Essex) cut with the same enzymes, yielding plasmid pXZ10. In this way, a C-terminal His-Tag is added to CslA, which expressing CslA with a 6-histidine tag at the C-terminus.

The *glxA* gene (SLI3188) was amplified without the sequence corresponding to its N-terminal cytosolic sequence (i.e. amino acids 1-11), using primers 3188-F/3188-R. The amplified fragment, cut with NcoI/HindIII, was then cloned behind a *pelB* signal sequence into pET22b (Novagen) cut with the same enzymes, yielding plasmid pXZ11. In this way, the expressed GlxA contains an additional Met-Gly dipeptide at the N-terminus after cleavage of the PelB leader and carries a His-Tag at the C-terminus.

To co-express CslA and GlxA, both genes were inserted into the pETDueT-1 plasmid (Novagen) in two steps. First, the *pelB-glxA* fragment was amplified from plasmid pXZ11 with primers pglxA-F/pglxA-R and ligated into pETDueT-1 as a NdeI/PacI fragment, yielding pXZ12. In this way, GlxA is produced and secreted in the periplasm without any tags. Secondly, primers cslA-F and cslA-R (containing a sequence for 12-histidine residues, see Table S2) were used to amplify *cslA* from plasmid pXZ10, after which the fragment was ligated into plasmid pXZ12 using XbaI and EcoRI, yielding plasmid pXZ13. To create pXZ44 (for co-expressing CslA and GlxA^12-261^) and pXZ45 (for co-expressing CslA and GlxA^12-54^), PCR-directed mutagenesis was used with pXZ13 and primer combinations of Flag-F/GlxA261-R and Flag-F/GlxA54-R, respectively. All plasmids were introduced into *E. coli* BL21(DE3) competent cells using heat-shock transformation (31).

### Membrane topology analysis of GlxA

To study the transmembrane topology of GlxA, the β-lactamase*-*encoding gene *blaM* without its signal sequence was fused to the 3’ end of *glxA*, as described previously (32). Only if BlaM is secreted, cells will be resistant to ampicillin. To this end, a *blaM* variant without its native signal sequence for secretion (*blaM_NS_*) was amplified from plasmid pHJL401 (33) with primers blaM-F/blaM-R. In parallel, full length *blaM* (including the region for the signal sequence, hereinafter referred to as *blaM_FL_*) was amplified from the same plasmid using the blaM_FL_-F/blaM-R primers. Meanwhile, nucleotides 1-96 of *glxA* or its full-length sequence (*glxA*_FL_) were amplified from *S. lividans* genomic DNA using primers glxA_1-96_-F/glxA_1-96_-R and glxA_blam_-F/glxA_blam_-R, respectively. Subsequently, the amplified products were cut using restriction enzymes NdeI-HindIII (*gxlA*_1-96_ and *glxA*_FL_), HindIII-EcoRI (*blaM*_NS_) and NdeI-EcoRI (*blaM*_FL_), and digested combinations of *gxlA*_1-96_ /*blaM_NS_*, *gxlA*_FL_/*blaM*_NS_ and *blaM*_FL_ were separately ligated into pXZ2 (17) that was cut with NdeI-EcoRI, yielding pXZ42, pXZ43 and pXZ36 (see Table S2), which in turn were used to express GlxA1-32-BlaMNS, GlxAFL-BlaMNS and BlaMFL, respectively.

*E. coli* DH5α harboring plasmid pXZ42, pXZ43 and pXZ36 were used to assess the membrane topology of GlxA, which were performed as described previously (32).

### Protein expression and purification

For protein expression, transformed colonies were randomly picked and inoculated in 250 ml Terrific Broth (TB) medium in a 2 liter flask. The cells were grown at 37°C while shaking until the OD_600_ was 0.4. Then, IPTG was added to the flasks (0.2 mM for the expression of CslA or GlxA individually, or 0.5 mM for co-expression of CslA with GlxA, GlxA^12-261^ or GlxA^12-54^) which were then further incubated at 16°C. After 18 hours induction, the cells were collected by centrifugation, resuspended in RB1 buffer (20 mM sodium phosphate buffer pH 7.2, 0.3 M NaCl, 10% glycerol) and then lysed by sonication (BANDELIN SONOOULS, power setting 30%, 2 seconds on/5 seconds off, 10 min duration). The crude membranes were collected by centrifugation for 60 min at 100,000 *g* in a Beckman Coulter Beckman Coulter Optima XE Ultracentrifuge equipped with a Beckman Ti45 rotor and immediately used or stored at −80°C until further use. The collected membranes were solubilized for 2 hours at 4°C in RB2 buffer (RB1 buffer supplemented with 1% n-Dodecyl-B-D-Maltoside (DDM)). The insoluble membrane fraction was separated from the soluble fraction by centrifugation at 100,000 *g* for 60 min. Then, the membrane fraction was incubated overnight at 4°C with 2 ml Co^2+^-resin (for the purification of CslA or GlxA monomers) equilibrated with RB3 buffer (RB1 buffer containing 0.05 % DDM). The resin was packed in a gravity flow chromatography column and washed with 50 ml WB1 buffer (RB1 buffer containing 25 Mm imidazole and 0.05 % Triton X-100). Proteins was eluted with 15 ml EB buffer (RB1 buffer containing 250 Mm imidazole and 0.05 % Triton X-100). The purification of the protein complexes (following co-expression of both proteins) was similar to the procedure for purifying CslA or GlxA alone, with the difference that 5 ml Ni-NTA resin was used and that after incubation with the membrane extract the resin was washed with 50 ml WB2 buffer (RB1 buffer containing 40 Mm imidazole and 0.05 % Triton X-100).

The eluted proteins were concentrated by centrifugation at 4000 rpm min^-1^ until the desired volume was reached and subsequently purified by size exclusion chromatography (SEC). Therefore, the Superdex 200 Increase 10/300 GL column (GE Healthcare) was equilibrated with RB buffer containing 20 mM sodium phosphate buffer pH 7.2, 0.2 M NaCl, 0.05 % Triton X-100. After collection, the eluted fractions were analysed using SDS-PAGE and concentrated using 10-kDa (for CslA or GlxA monomers) or 100-kDa (for protein complexes) cut-off centrifugal filters (Millipore).

### SDS-PAGE and immunoblotting assay

SDS-PAGE was performed with the Bio-Rad Mini-PROTEIN system using 7.5% gels (34). To visualize protein bands, gels were stained with Coomassie Brilliant Blue (Sigma). The authenticity of the proteins was verified by immunoblotting, which was essentially performed as described (18). For the detection of His-tagged CslA, a His-Tag monoclonal antibody (Proteintech) and an Anti-Mouse lgG-Alkaline Phosphatase (Sigma) were used as the primary and secondary antibody, respectively. The antibodies used for the detection of GlxA were identical to those described before (18). For the detection of 3×Flag-tagged GlxA truncations, a Flag-Tag monoclonal antibody (Proteintech, 1:1,000 dilution) and an Anti-Mouse lgG-Alkaline Phosphatase (Sigma, 1:5,000 dilution) were used as the primary and secondary antibody, respectively. Images were collected using an Epson Perfection V37 scanner.

### In vitro synthesis of glucans

For the in vitro synthesis of glucans, the reaction mixtures contained 0.1 or 0.05 mg ml^-1^ protein(s), 20 mM sodium phosphate, Ph 7.2, 0.1 M NaCl, 5 mM cellobiose, 0.05 % triton X-100, 5 mM UDP-glucose (UDP-Glc, Promega), 20 mM MgCl_2_ and 30 mΜ cyclic-di-GMP (Sigma). All reactions were set up as below. 50 μl of solution A, (20 Mm sodium phosphate, pH 7.2, 10 mM cellobiose, 40 mM MgCl_2_ and 60 mΜ cyclic-di-GMP) (23), 0.05% triton X-100 and 10 mM UDP-sugars, was placed into two Eppendorf tubes. To one of the tubes, 50 μl of 0.2 or 1 mg ml^-1^ protein in RB buffer was added, while 50 μl of RB buffer was added to the other tube, serving as the enzyme blank control. For detecting if cyclic-di-GMP is required for the activity of the CslA-GlxA complex, this compound was removed from the reaction setup without any further changes. Following incubation at 37°C for 60 min under shaking conditions (250 rpm), the mixtures were centrifuged at 16,000 *g* for 20 min to separate the (soluble) released UDP from the insoluble glucans. These separated fractions were subsequently used for glycosyltransferase activity measurements and glucan constitution analysis (see below), respectively.

### Quantitative measurement of the glycosyltransferase activity of CslA, GlxA or CslA-GlxA

The glycosyltransferase activity of the CslA-GlxA complex was quantitatively measured by using the UDP-Glo^TM^ glycosyltransferase assay kit (Promega). This assay detects the content of free UDP, which is released during the glycosyltransferase reaction. After separation of the insoluble glucans by centrifugation, 15 μl of the UDP-containing supernatant was mixed with 15 μl of freshly prepared UDP-Glo^TM^ Detection Reagent, which also terminates further glucan synthesis. After 1 h of incubation at room temperature in the dark, luminescence was measured using a TECAN Spark M10 plate reader. All values were corrected using the enzyme blank controls, and the standard curve line of luminescence-UDP was used to convert measured luminescence to the UDP concentration. All measurements were performed in triplicate.

### Analysis of the in vitro synthesized glucans

To characterize the glucans produced *in vitro* by the CslA-GlxA complex, linkage analysis was performed following enzymatic digestion. Briefly, the glucans from the *in vitro* reactions were collected by centrifugation, washed with Milli-Q water, and resuspended in 200 μl enzyme solution (25 mM Tris, Ph 7.5, 100 mM NaCl) containing 20 μg ml^-1^ cellulase (Sigma) or 100 μg ml^-1^ endo-β-1,3 glucanase from *Trichoderma* sp. (Megazyme). Enzymatic degradation was performed at 37°C for 48 h under shaking conditions (250 rpm min^-1^), after which the reaction was centrifuged at 16,000 g for 20 min. Reducing sugars in the supernatant were measured using DNS reagent (35). A standard curve line (made with a glucose standard (Sigma)) was used to convert the absorbance at 590 nm to the concentration of reducing sugars. All measurements, performed in triplicate, were corrected using the enzyme blank control.

To further characterize the glucans, the oligosaccharides in the obtained cellulase digests were subsequently analyzed using Matrix-Assisted Laser Desorption/Ionization Fourier Transform Ion Cyclotron Resonance Mass Spectrometry (MALDI FT-ICR MS) (36, 37). Approximately 300 μg ml^-1^ of reducing oligosaccharides present in the supernatant fraction after cellulase digestion were 10-fold diluted with a 50:49.95:0.05 (v/v/v) acetonitrile:water:formic acid solution. Then, 1 µl of each diluted sample was spotted on MALDI 96 Ground Steel target plates with 1 µl of 1 mM NaCl and 1 µl of a 10 mg ml^-1^ “super-DHB” (a 9:1 (w/w) mixture of 2,5-dihydroxybenzoic acid and 2-hydroxy-5-methoxybenzoic acid; Sigma) solution in 50:50 (v/v) acetonitrile:water. The spots were left to dry at room temperature. All mass measurements were performed on a 15 T solariX XR Fourier transform ion cyclotron resonance (FT-ICR) mass spectrometer (Bruker Daltonics, Bremen, Germany) equipped with a CombiSource and a ParaCell. A Smartbeam-II laser system (Bruker Daltonics) was used to perform MALDI measurements using a laser frequency of 500 Hz and 200 laser shots per scan. MS1 measurements were performed in the *m/z*-ranges of 1,011-8,000 and 2,000-30,000. Collision-induced dissociation measurements were performed on selected precursor ions using a collision energy of 50 V and detection of CID fragment ions in the *m/z*-range 92-8000.

### Statistical analysis

GraphPad Prism software (version 8.0.2) was used for statistical analyses. Paired t-tests were used to calculate the significance in pairwise comparisons.

### Bioinformatics analyses

All amino acids sequences of cellulose synthases or cellulose-like synthases were downloaded from UniProt (https://www.uniprot.org/). Phylogenetic tree construction was performed using MEGA 7 (38). The alignment of amino acids sequence was performed Espript 3.0 (https://espript.ibcp.fr/ESPript/ESPript/). Structure prediction of CslA (aa. 44-599)-GlxA (aa. 12-645) complex were performed with Alphafold 2.0 (39, 40).

## Data availability

All data are contained within the manuscript.

## Supporting information

This article contains supporting information.

## Acknowledgements

We thank Dr. Patrick Voskamp from the Leiden Institute of Chemistry and Dr. Lennart Schada von Borzyskowski from the Institute of Biology for their help with the size exclusion chromatography. The work was supported by a Vici grant from the Dutch Research Council (I) to D.C. (VI.C.192.002) and a CSC scholarship to X.Z.

## Abbreviations

The abbreviations used are:

GT: glycosyltransferase
CslA: cellulose synthase-like protein A
GlxA: galactose oxidase-like protein A
CslZ: cellulose synthase-like protein Z
LpmP: lytic polysaccharides monooxygenase for peptidoglycan
CesAs: cellulose synthases
Csl: cellulose synthase-like
BcsA: bacterial cellulose synthase A
BcsB: bacterial cellulose synthase B
CcsA: clostridial cellulose synthase A
CcsB: clostridial cellulose synthase B
MALDI FT-ICR MS: matrix-assisted laser desorption/ionization fourier transform ion cyclotron resonance mass spectrometry
Glc: glucose
UDP: uridine diphosphate
TM: transmembrane
PG: peptidoglycan
c-di-GMP: 3′,5′-cyclic diguanylic aci
aa.: amino acids

